# Pilot MRI study of carbon monoxide (CO) against ischemic stroke in mice: blood brain barrier integrity and metabolic pattern

**DOI:** 10.1101/2022.11.10.515956

**Authors:** Sara R. Oliveira, João Castelhano, José Sereno, Lorena Petrella, Miguel Castelo-Branco, Helena L. A. Vieira, Carlos B. Duarte

## Abstract

Although stroke is the main cause of brain damage worldwide, stroke therapies are based on blood reperfusion and do not target cerebral parenchyma. Ischemic stroke (representing 87% of all strokes) causes cerebral damage due to oxygen and tissue energy depletion, which lead to acidosis, inflammation, excitotoxicity and oxidative stress. Carbon monoxide (CO) is an endogenous gasotransmitter produced by heme oxygenase cleavage of the heme group. CO promotes cytoprotection by limiting inflammation and preventing cell death in several tissues including the brain. Previous studies have demonstrated the protective role of CO in the mouse ischemic stroke model, middle cerebral artery occlusion (MCAo) by histological analysis when CO is when applied before ischemia. Herein, there are two main novelties. First CO is administrated following stroke, which better mimics its potential future use as therapeutic drug. Secondly, imaging techniques were used to elucidate the effect of this gasotransmitter at the metabolic, vascular and anatomic levels.

The putative neuroprotective effects of CO following MCAo were assessed by 3 i.p. injections of the CO-releasing molecule CORM-A1 (3 mg/kg), administered 6, 24 and 48h after reperfusion. Magnetic Resonance Imaging was performed 1 day and 7 days after reperfusion using T2-weighted, diffusion weighted images, proton spectroscopy (^1^H-MRS) and perfusion (dynamic contrast enhanced images). ^1^H-MRS also allowed the comparison between metabolite signatures at day 1 versus 7 day following MCAo. Furthermore, CORM-A1 limited the loss of blood-brain barrier (BBB) integrity as it reduced the edema formation. Furthermore, the CO donor minimized the metabolite load loss at an early stage after MCAo, both in striatum and cortex.

In conclusion and based on MRI analysis, CO has a protective role in the recovery from stroke injury, mainly by acting on BBB integrity and brain metabolism.

## INTRODUCTION

Carbon monoxide (CO) is a gasotransmitter endogenously produced in many organs. The brain is one of the tissues with the highest activity of heme oxygenase, the enzyme that produces CO (Maines 2000). A wide range of physiological roles are played by CO in the brain, including cytoprotective effects such as anti-apoptotic and anti-inflammatory, metabolic improvement, vasomodulation, and promotion of neurogenesis (Queiroga et al. 2015; Almeida et al. 2016, 2018; Oliveira et al. 2019; Figueiredo-Pereira et al. 2020). Neuroprotective actions of CO were described in *in vivo* models of epilepsy (Parfenova et al. 2012a), of adult permanent ischemic stroke (Wang et al. 2011), of hemorrhagic stroke (Yabluchanskiy et al. 2012) and in perinatal ischemia (Queiroga et al. 2012), among others. However, the mechanisms underlying the neuroprotective effects of CO are still poorly understood.

Ischemic brain injury is a leading cause of mortality and morbidity in western countries. Tissue plasminogen activator (tPA) and thrombectomy are the only approved therapies for acute ischemic stroke (Wardlaw et al. 2014; Meurer et al. 2016). However, these therapeutic strategies can only be used in short time window and do not target brain tissue (Schwamm et al. 2013). Despite the failure of the previous generation of drugs, new approaches have been pursued for stroke therapy, namely the stimulation of endogenous cytoprotective mechanisms. To overcome the complexity of stroke, one may hypothesize that a molecule that targets as many stroke hallmarks as possible can be a promising therapeutic candidate. Studies have been performed to evaluate the therapeutic potential of CO in stroke models, mainly by administration of CO in a prophylactic manner. A few studies have tested the effect of CO administered after injury at different time points (from several days before injury to 3 days after reperfusion) (Zeynalov and Dore 2009; Klaus et al. 2010; Wang et al. 2011; Queiroga et al. 2012). A drawback in most of the studies described above is the choice of the time point of CO administration; treatment with CO before the ischemic insult has a limited clinical relevance. Moreover, since the protective readouts were often collected soon after injury, the long-term effects remain uncharacterized.

In the present study, we investigated the cerebral effects resulting from systemic administration of low doses of CO after brain ischemia, using a mice model of transient middle cerebral artery occlusion (tMCAo). This approach evaluates the therapeutic potential of CO in a clinically relevant scenario: the effect of the gasotransmitter was tested at time periods longer than 4.5 h after the ischemic injury as an effort to overcome the limitation of tPA’s short therapeutic window. The route of CO administration, i.p. injection, was also chosen to be easily translated into the clinical practice. As CO presents a complex biology, the effects of this gasotransmitter were evaluated in an integrated manner, characterizing the effects on the vasculature and metabolism in a longitudinal study. We found that 3 intraperitoneal CO injections given within 6h to 48h after the stroke onset promote a robust reduction in BBB permeability. Moreover, CO-treated mice subjected to tMCAo presented a tendency to a reduction in infarct volume measured by MRI at day 1, which was correlated with a limited metabolite loss, in particular in the cortical region.

## MATERIALS AND METHODS

### Experimental procedure

The mice were randomly divided into four groups (n=5/6 *per* group): i) Sham group, mice were treated with saline; ii) Sham treated with CORM-A1; iii) tMCAo group, mice were treated with saline and iv) tMCAo treated with CORM-A1. The treated groups received CORM-A1 via i.p. injection (3 mg/kg in saline) at 6 h, 1 day and 2 days after MCAo. Animal experiments were conducted according to the European Council Directives on Animal Care and were reviewed and approved by DGAV, Portugal.

### Transient middle cerebral artery occlusion (tMCAo)

Surgical procedures were performed according to the protocol published as the latest Standard Operating Procedure (SOP) (Prinz et al. 2010). The national and local animal welfare authorities approved all animal experiments. Male C57BL/6 mice were obtained from Charles River at the age of 7-8 weeks and were used in the experiments at the age of 8–10 weeks. All animals had access to food and water *ad libitum*, and were kept under a 12 h light/dark cycle. No specific exclusion criteria were set.

The right middle cerebral artery (MCA) was occluded for 45 min. In summary, mice were initially anesthetized by inhalation of 2.5% isoflurane in O_2_:N_2_O (30:70). Thereafter, mice were placed on a heating pad and anesthesia was subsequently reduced and maintained at 1.5-1.8% (using an open mask). A rectal temperature probe was used to monitor body temperature, which was kept at 37°C (see Figure S2). An optical fiber probe (Probe 318-I, Perimed, Sweden) was firmly attached to the skull and connected to a laser doppler flow meter (Periflux System 5000, Perimed), to monitor changes in regional cerebral blood flow (rCBF). A 6-0 silicon-coated (about 9-10mm is coated silicon) monofilament suture (602334PK10, Doccol Corporation) was introduced into the ECA (external carotid artery), which was monitored by a sudden drop in rCBF (Supplementary Data Figure S2). Bupivacaine (0.150 μL, 0.05%, Marcain™, AstraZeneca, Sweden) was injected around the wound to reduce pain. In order to avoid *post*-surgical hypothermia, animals were placed in an incubator at 35°C for the first 2 h after the procedure and then transferred to an incubator at 33°C (overnight). 30 min we administered 0.5 mL of 5% glucose in saline subcutaneously. As animals were allowed to recover for 7 days, we further administered 0.5 mL of 5% glucose subcutaneously daily, up to day 4 *post*-surgery (which is when weight loss ceases). Body weight was controlled daily up to the experimental endpoint. In sham-surgeries, the filament was advanced up to the internal carotid artery, and was withdrawn before reaching the MCA.

### CO administration

CORM-A1, a CO releasing molecule (Motterlini et al. 2005), was used to systemically deliver CO in mice. The CORM-A1 (Sigma) stock solution was prepared in 0.9% NaCl solution and stored at −20°C to avoid loss of released CO. 3 mg/kg of CORM-A1 were administered i.p at 6 h ± 0.2 h, 32 h 39 min ± 5h and 56 h 08 min ± 6 h.

### Infarct volume quantification by MRI

Stroke volume was quantified by T2-weighted magnetic resonance imaging (T2*-MRI). For stroke volumetry, hyperintense areas of ischemic tissue in T2-weighted images were assigned with a region of interest tool. This enabled a threshold-based segmentation that was performed by connecting all pixels within a specified threshold range around the selected seed pixel and resulted in a 3D object map of the whole stroke region. The total volume of the whole object map was calculated automatically. For quantification of the regional ischemic lesion volumes, axial MR images were divided into three equal slices from medial to lateral with NIH ImageJ. The volume of hyperintense areas in each brain section was calculated with the Analyze 5.0 software. Brain parenchyma was cropped for presentation purposes.

### Cerebral perfusion quantification by MRI

Mice were anesthetized with 1.5% isoflurane in air, and placed on controlled temperature beds through water baths (Haake SC 100, Thermo Scientific, USA) with tooth bar and head restraint to reduce motion artifact. Body temperature was maintained at 37ºC with assessment of cardiorespiratory function (1030, SA Instruments Inc., NY, USA). MRI was performed on a 9.4T MR pre-clinical scanner (Bruker Biospec, Billerica MA, USA) equipped with a standard Bruker crosscoil setup using a volume coil for excitation (86/112 mm of inner/outer diameter) and quadrature mouse surface coil for detection (Bruker Biospin, Ettlingen, Germany). The system was interfaced to a Linux PC running Topspin 3.1PV and Paravision 6.0 (Bruker Biospin). High resolution morphological images were acquired with a 2D T2-weighted turbo RARE (rapid acquisition with relaxation enhancement) sequence with TR = 3.6 ms, TE = 33.0 ms, matrix 256 × 256 and field of view 20.0 × 20.0 mm reaching an in plane spatial resolution of 78*78 µm^2^, rare factor 8, axial slice thickness 0.5 mm and 5 averages resulting in 9 min 36 s of scan time.

Blood brain barrier permeability was evaluated after injection of Gadovist^®^ (0.1 mmol/kg) using the dynamic contrast-enhanced magnetic resonance imaging technique. The following parameters were used in 2D DCE-FLASH: TE/TR = 2.5/191.145 ms, flip angel = 70º, FOV= 20 × 20 mm, matrix= 156 × 85, slice thickness 0.5 mm, number of slices= 18, number of repetitions= 40 and time acquisition of 1 h 3 min 20 s.

### ^1^H-MRS

Data were collected on a volume of interest selected in accordance with coronal, sagittal and axial T2-weighted images, and adjusted to fit the anatomical structure of interest and to minimize partial volume effects. The B0 map was acquired and MAPSHIM was employed to automatically adjust first and second-order shim coils, with iterative correction. Spectral line widths of water around 13–18 Hz were obtained. A PRESS sequence was used in combination with outer volume suppression and VAPOR water suppression. The following parameters were used: TR = 2500 ms, TE = 16.225 ms, number of averages = 720, number of acquired points = 2048, yielding a spectral resolution of 1.22 Hz/point. For each animal, an unsuppressed water signal (TE = 16.225 ms, TR = 2500 ms, 16 averages, scanning time = 40 sec and none water suppression) was acquired immediately before acquiring the water-suppressed spectrum.

Data were saved as Free Induction Decays (FIDs) and corrected for the frequency drift and for residual eddy current effects using the reference water signal. The ^1^H NMR peak concentrations for major metabolites (*e*.*g*., *N*-acetylaspartate (NAA), γ-aminobutyric acid (GABA), taurine, glutamine and glutamate (Glx)) was then analyzed using the LCModel software package (Stephen Provencher Inc., Oakville, Canada; Provencher 1993), and the results were given in relation to the water content in the tissue. Briefly, the LCModel analysis calculates the best fit to the acquired spectrum as a linear combination of a model based on a set of brain metabolites from the LCModel basis set. The Cramer-Rao lower bound provided by LCModel was used as a measure of the reliability, and metabolite concentrations with Cramer-Rao lower bound higher than 24% were not included in the analysis.

### Statistical Analysis

Two-way repeated-measures (RM) ANOVA with Bonferroni’s *post hoc* tests for comparison of sample treatment effects within Sham or tMCAo was used. Two-way RM ANOVA, followed by Tukey’s *post hoc* analysis, was performed for statistical analysis of the results obtained in the MRI stroke volume assessment at 1 and 7 days after tMCAo.

### Methods to prevent bias

In summary, out of the 52 animals used, 2 were excluded due to a hemorrhage during surgery and 2 were excluded as they died during MRI acquisition. 48 animals were included in the study from which 21 reached the end point (Supplementary Data, Figure S1).

Animals were randomized before the experiments by a scientist who was not involved in the tMCAo experiments, in the administration of CO or in the behavioral tests. MRI and behavior data analysis were performed blind for the grouping, and the treatment allocation was secret. MRI data acquisition was also performed by scientists who were not involved in the surgery experiments or CO administration, and both the grouping and the treatment allocation were concealed.

## RESULTS

### CO reduces BBB leakiness

The putative effects of CO in neuroprotection were evaluated by i.p. injection of the donor CORM-A1 (3 mg/kg) 6 h after tMCAo (45 min occlusion), followed by administration of the same dose 1 and 2 days later. The use of this concentration of CORM-A1 ensures that carboxyhemoglobin was not significantly increased compared to baseline (Csongradi 2012), thus avoiding any toxicity.

Blood-brain barrier (BBB) leakiness plays a major role in neuronal damage after transient brain ischemia, enhancing tissue damage (Krueger et al. 2015; Kassner and Merali 2015). There are multiple evidence for modulatory and protective effects of CO on the vasculature (Knecht et al. 2010; Leffler et al. 2011). Therefore, we hypothesized that the CORM-A1 could improve BBB integrity. To address this question, a MRI-based technique was used. A gadolinium-based contrast agent was given i.p. and its signal was quantified in brain tissue (Jiang et al. 2005; Wunder et al. 2012). Since under normal physiological conditions the BBB excludes the contrast agent, any increase in signal intensity is interpreted as BBB breakdown. The results of Figure 1 show a significant increase in the permeability of the BBB when evaluated 1 day or 7 days after tMCAo. Under the latter conditions, CORM-A1 administration significantly reduced BBB damage in mice subjected to tMCAo (Figure 1B), which was very clear when the cumulative signal of the contrast agent was represented with a color code (Figure 1A). The total accumulation of the contrast agent at day 7 in animals exposed to focal ischemic injury followed by administration of CORM-A1 was similar to that observed after tMCAo at day 1, suggesting that CO prevents the delayed increase of BBB permeability, rather than the initial alterations. The untreated MCAo cohort showed a 3-fold increase in the accumulation of contrast agent in the same period of time (Figure 1B).

**Figure 1.**
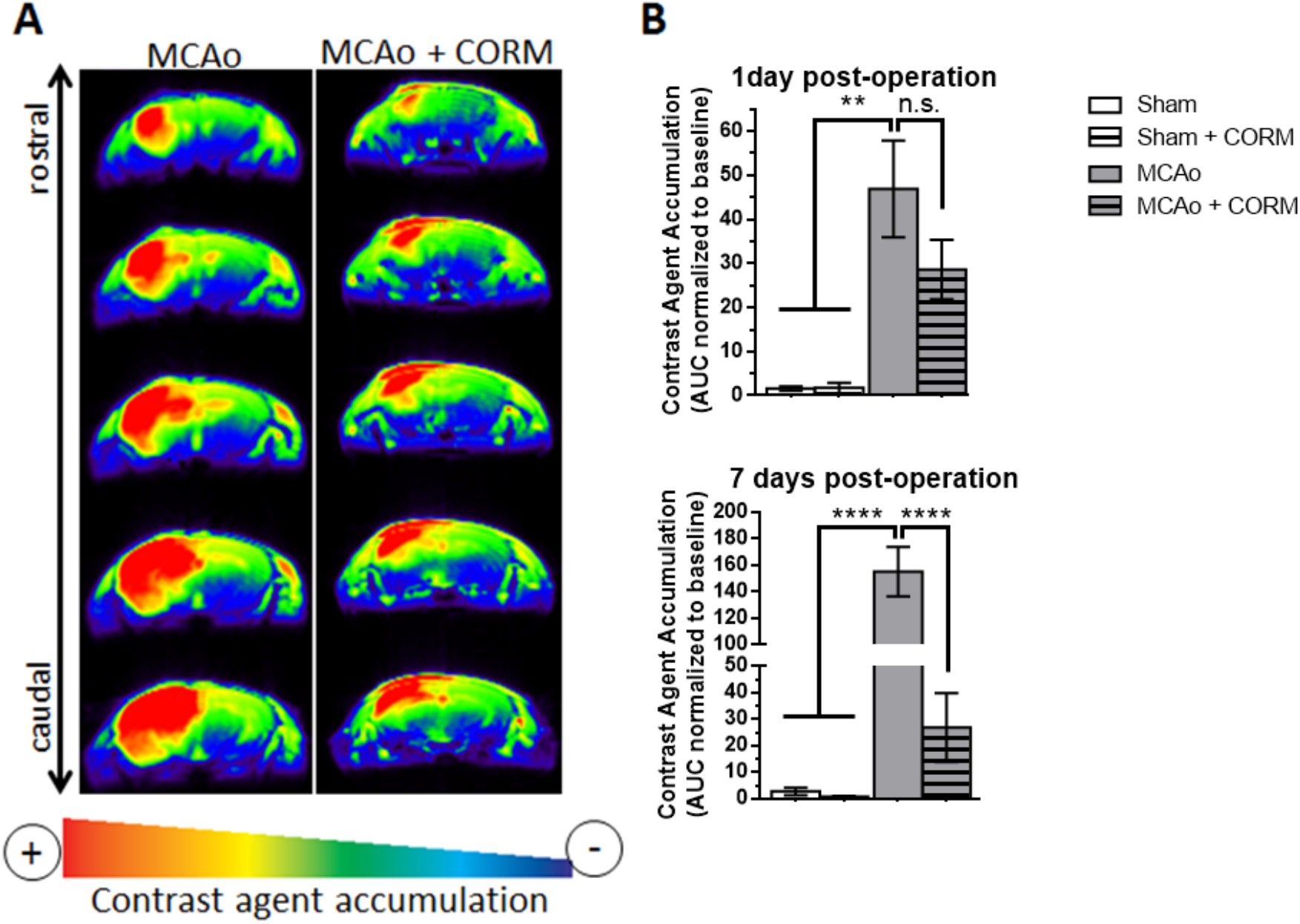
Alterations in the BBB permeability after tMCAo. Where indicated, the animals were treated with CORM-A1 (3 mg/kg) in 3 doses (6 h, 1 day and 2 days *post* injury). Accumulation of contrast agent in the brain tissue occurs in regions where the BBB in compromised. (A) Representative images showing accumulation of the contrast agent 7 days after tMCAo in mice. (B) Quantification of the signal intensity resulting from the accumulation of the contrast agent. White bars represent sham-operated mice and grey bars represent MCAO-operated mice. Clear and striped bars represent control and CO-treated mice, respectively. \*\**p*<0.01, \*\*\*\**p*<0.0001, n.s. – not significant, determined with unpaired Student’s *t*-test.

Taken together, the results show the prolonged effects of CO in the protection of the BBB after ischemic injury, which resulted in preservation of the BBB integrity.

### CO limits infarct lesion 24h after MCAo

To determine whether the preservation of the BBB integrity in CO treated mice correlates with a smaller infarcted area, MRI was used to quantify the ischemic brain tissue:. When total lesion volume was evaluated by MRI, CO treated animals presented a slightly smaller infarct size at day 1 compared to the untreated group (Figure 2A), but the results were not statistically significant. Since the cerebral cortex and striatum are the most affected brain regions in the MCAo model of transient focal ischemia (McColl et al. 2004), the ischemic injury was also specifically quantified in these two areas. This categorization showed that the cortical lesion, but not the striatal injury, was slightly reduced when the animals were treated with CORM-A1 after MCAo, but the results were not statistically significant (Figure 2B, C). Analysis of the lesion volume at day 7 after MCAo showed no effect of CORM-A1 administration.

**Figure 2.**
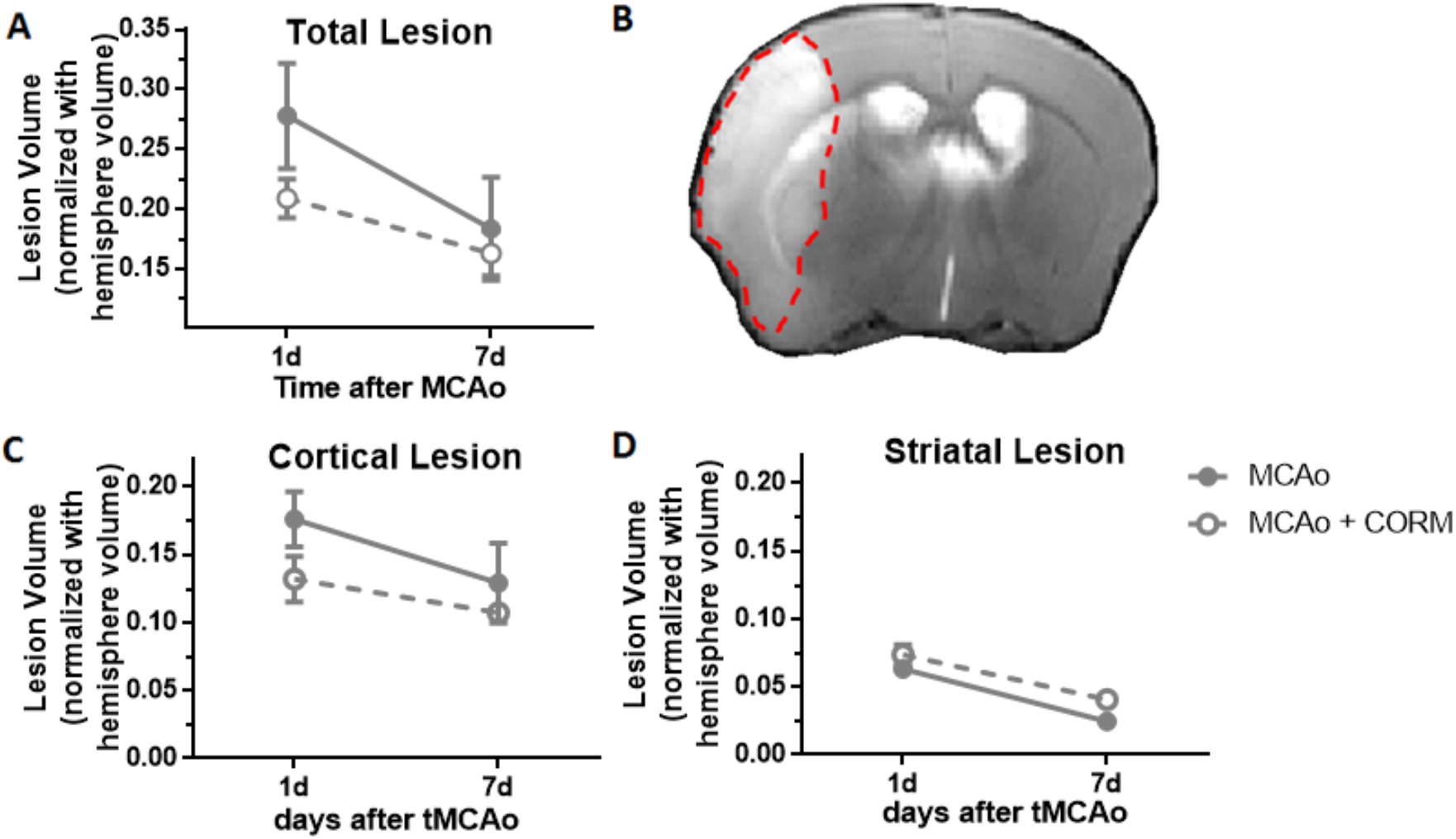
MRI quantification of infarcted brain volume in mice subjected to tMCAo. Where indicated, the animals were treated with CORM-A1 (3 mg/kg) in 3 doses (6 h, 1 day and 2 days *post* injury). Anatomic volumetries were measured at day 7. The results obtained in untreated mice subjected to MCAo are represented by filled circles, while empty circles showed the results obtained in animals subjected to the ischemic insult followed by CORM-A1 treatment. (A) Total lesion volume; (B) representative image of the infarcted mouse brain with the hyperintense lesioned tissue limited with a dashed red line; (C) lesion volume affecting the cerebral cortex; (D) lesion volume affecting the striatum. The results are the average ± SEM n = 5. Statistical analysis was performed using the two-way ANOVA followed by the Bonferroni *post*-test.

In summary, the CO dosage given was insufficient to decrease the infarct volume in mice subjected to tMCAo, specially at day 7 after the lesion. Although a slight protection was observed in the cerebral cortex at day 1 *post*-MCAo, the effect was not statistically significant and was not observed at a later point.

### CO minimizes cortical metabolic loss 24h after tMCAo

To further characterize the differential effects of CO when tested at day 1 and day 7 after tMCAo (*e*.*g*. in the BBB permeability), metabolic profiles (Figure 3A) were obtained at both time points. ^1^H-MRS was applied in a voxel placed in the cerebral cortex (Figure 3B) to determine whether CORM-A1 administration prevents the ischemia-induced metabolic changes, as previously describe in other contexts (Ryter and Choi 2009; Mahan 2012; Almeida et al. 2012). Nine metabolites were analyzed by this method (named here as metabolic load) in order to have a general assessment of the alterations in brain metabolism: alanine, phosphocholine, phosphocreatine, lactate, taurine, *myo*-inositol, gluthathione, glutamate, glutamine, GABA, *N*-acetylaspartate. tMCAo reduced the metabolite load in the ipsilateral hemisphere, when assessed 1 day after the ischemic injury, while no effect was observed in the contralateral hemisphere. Administration of CORM-A1 significantly reduced the metabolic changes in the ipsilateral hemisphere at this time point, to levels similar to the sham cohorts (Figure 3C), in contrast with the results obtained at day 7, when CO had no effect on MCAo-evoked metabolite loss (Figure 3D). Under the latter conditions both MCAo subjected groups presented the same metabolite load loss.

**Figure 3.**
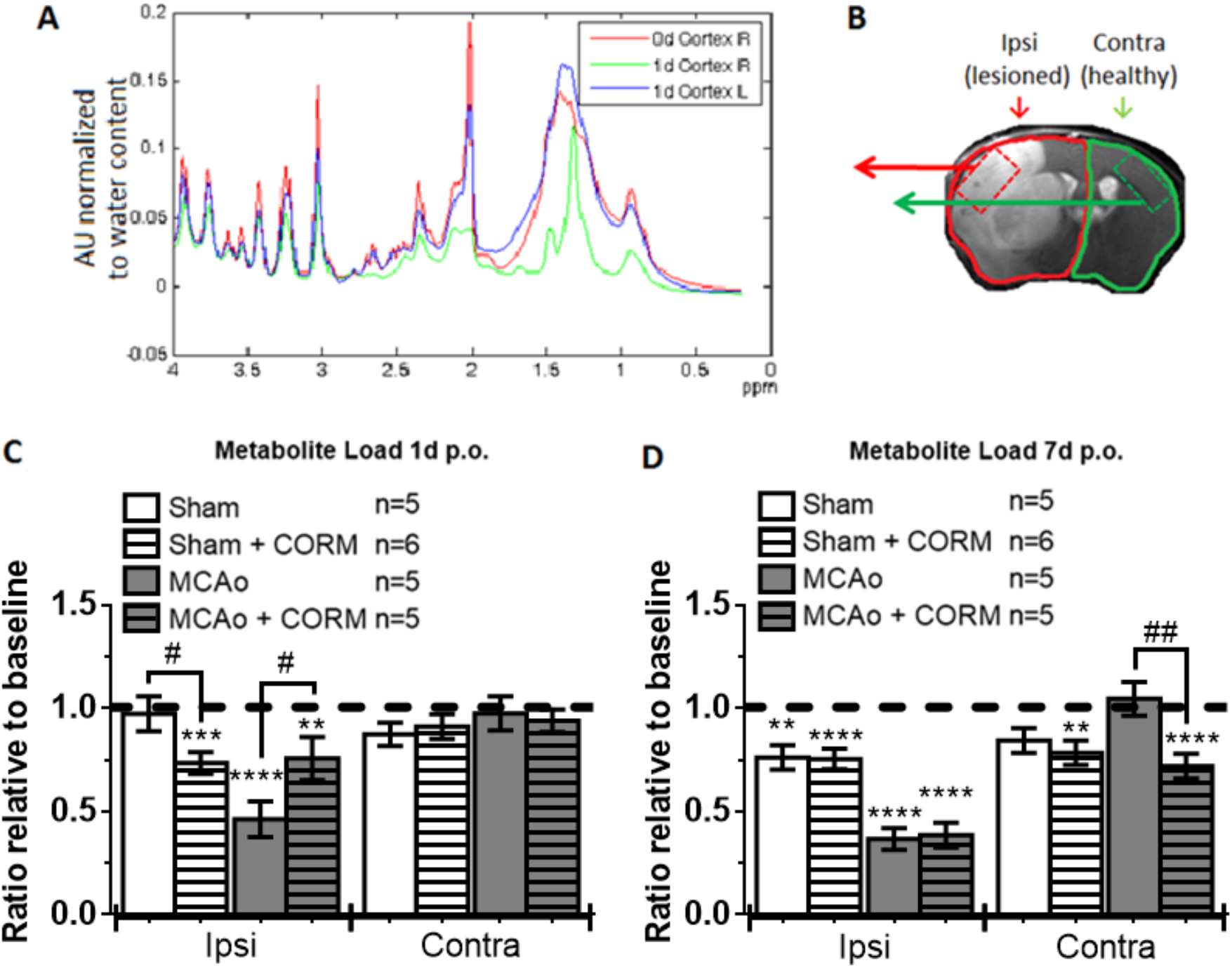
Modulation of the MCAo-induced alterations in the metabolic profile in the cerebral cortex by CORM-A1. Metabolite load includes the following metabolites: alanine, phosphocholine, phosphocreatine, lactate, taurine, *myo*-inositol, glutathione, glutamate, glutamine, GABA and *N*-acetylaspartate. Where indicated, the animals were treated with CORM-A1 (3 mg/kg) in 3 doses (6 h, 1 day and 2 days *post* injury). (A) Representative plot of distribution of metabolites in control conditions and at days 1 and 7 after MCAo, as determined by ^1^H-MRS quantification. (B) Voxel placement in the ipsi- and contralateral hemispheres. (C) Ratio of metabolite load at day 1 after MCAo. (D) Ratio of metabolite load at day 7 after MCAo. White bars represent sham-operated mice and grey bars represent MCAO-operated mice. White and striped bars show the results obtained in control conditions and in mice treated with CORM-A1, respectively. \*\**p*<0.01, \*\*\**p*<0.001, \*\*\*\**p*<0.0001 calculated with one-way ANOVA with Bonferroni *post*-test calculated as a percentage of the control, determined before the surgery (mean ± SEM). # *p*<0.05 determined using the unpaired Student’s *t*-test.

These data clearly indicate that CO treatment decreases the early (1 day) metabolic changes that occur in the cerebral cortex after tMCAo. However, this was a transient effect since no effect was observed when the metabolic load was evaluated at day 7 after tMCAo.

### CO administration is beneficial for the maintenance of the metabolite load in the neuronal and astrocytic populations in the cerebral cortex

To determine whether CO treatment has a differential effect on the metabolism of neurons and astrocytes, a group of metabolites relevant for neuronal survival (*N*-acetylaspartate (NAA), glutamate (Glu) and taurine (Tau)) (Lei et al. 2009; Saransaari and Oja 2010) and a metabolite enriched in astrocytes (*myo*-inositol) (Harris et al. 2015) were evaluated in the cerebral cortex at day 1 after tMCAo. As proposed by others (Berthet et al. 2011), the composite score NAA + Glu + Tau predicts better the outcome of an ischemic insult than NAA or lactate alone. In the scatterplot representing glutamine *versus* the composite score, a good separation between Sham and MCAo subjected cohorts was obtained (Figure 4A). In accordance with the metabolic load and infarct volume at day 1, mice subjected to tMCAo followed by administration of CORM-A1 are more similar to the Sham cohorts than to untreated mice subjected to transient focal ischemia. Moreover, as this composite score is constituted by a group of metabolites relevant to the neuronal population, we may hypothesize that this score may account for an increased survival of the neuronal population.

**Figure 4.**
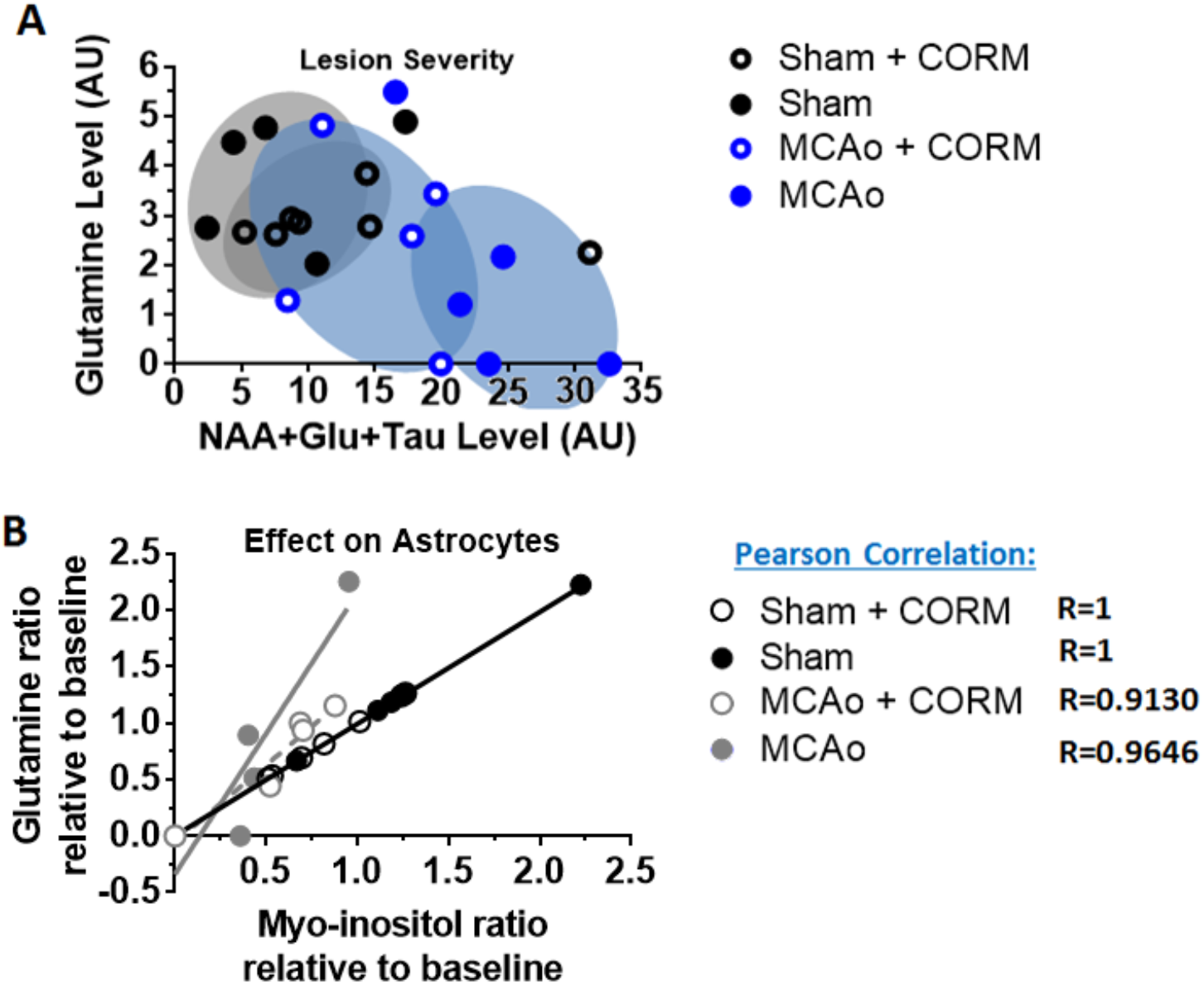
Severity of the lesion after tMCAo and astrocytic metabolic status in the cerebral cortex. Data were obtained from ^1^H-MRS at day 1 *post* tMCAo, and where indicated the animals were treated with CORM-A1 (3 mg/kg) in 3 doses (6 h, 1 day and 2 days *post* injury). (A) The distribution of NAA + Glu + Tau is plotted vs glutamine levels. The relation between the levels of two metabolites enriched in astrocytes, glutamine and *myo*-inositol, is plotted in panel (B).

When evaluating the correlation between the levels of *myo*-inositol and glutamine, two metabolites that are enriched in the astrocytic population, mice subjected to tMCAo followed by treatment with CORM-A1 presented a correlation very similar to sham treated groups (Figure 4B). In contrast, untreated mice with an ischemic lesion showed a distinct profile when compared with sham groups. In conclusion, administration of CO after the ischemic injury has a beneficial metabolic effect on both the neuronal and astrocytic populations, decreasing the severity of the lesion.

Taken together, these results reinforce the idea that CO acts in different cell populations, protecting both neurons and astrocytes, at least in part by maintaining the levels of key metabolites.

### CO ameliorates survival

Mortality reduction is also an important feature when evaluating the effect of putative therapeutic strategies for stroke. Administration of CORM-A1 after tMCAo, at 6 h, 1.3 d and 2.3 d after injury, during the therapeutic window, reduced the rate of mortality (Figure 5). However, the protective effects decreased after the last dose of CO (after day 3), and at day 7 following MCAo mice treated with CORM-A1 showed a survival rate similar to the non-treated controls (Figure 5). Survival data correlates very well with the lesion volume and metabolic load variations, since a certain degree of protection was observed during the therapeutic window (0-2 days after lesion).

**Figure 5.**
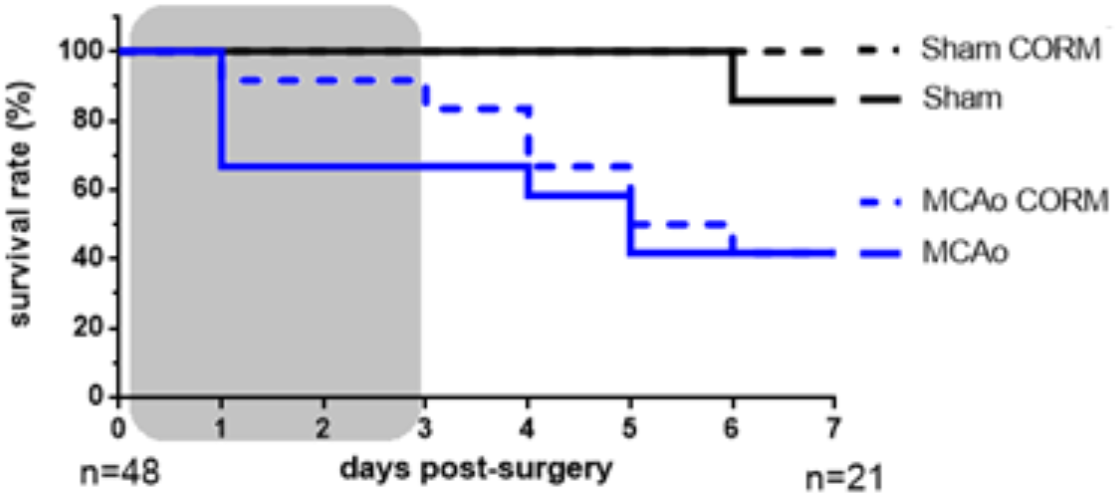
Effect of CO treatment on the survival rate of mice subjected to MCAo. CORM-A1 (3 mg/kg) was administrated in 3 doses (6 h, 1 day and 2 days *post* injury). The grey rectangle represents the therapeutic window applied. Black lines represent sham-operated mice and blue lines represent MCAO-operated mice. Full and dashed lines represent control and CO-treated mice, respectively.

In conclusion, the CO dose given has potential to improve outcome after ischemic stroke. However, the CO therapy needs to be optimized, including the timing of administration and the number of CO doses.

## DISCUSSION

In this integrative study, we investigated the putative neuroprotective effects of CO in a mouse stroke model. The results showed protective effects of CO at different levels, when administered from 6 h after injury: i) strong reduction of BBB leakage; ii) serial MRI data showed a tendency for smaller brain infarction in the cortical area of CO treated animals at day 1 *post* MCAo; iii) at day 7 *post* MCAo, CORM-A1 treatment did not affect the lesion volume as determined by MRI; iv) minimization of metabolite load loss in the cerebral cortex at day 1 after MCAo; v) CO reduced the mortality only when administered during the therapeutic window. Together, the evidence gathered in the current study indicates that CO has therapeutic potential as a *post*-stroke treatment.

BBB assessment several types of damage following stroke, with its cellular constituents (astrocyte end feet, pericytes, endothelial cells, smooth muscle cells and a variety of basement membranes) presenting different susceptibilities to the ischemic insult. The contribution of BBB for stroke pathogenesis has recently emerged as a focus for new therapeutic strategies (Kassner and Merali 2015). The results obtained in this work showed that CORM-A1 administration significantly prevents the early increase in BBB permeability after tMCAo, as evaluated 1 day after the lesion. The protective effects of CO on the integrity of the BBB function resemble the effects previously described for this gasotransmitter in several of its components, namely astrocytes (Queiroga et al. 2016; Choi et al. 2016b), pericytes (Choi et al. 2016a) and vasculature (Leffler et al. 2011). These observations were performed both in ischemic and non-ischemic conditions. Therefore, effects of CO in the stabilization of the BBB may arise from the combined effects on several of its components. However, it remains to be determined whether CO preserves the crosstalk between the different BBB components, acts through its anti-oxidant capacity and/or simply promotes vasodilatation. The latter effect of CO was reported in several other contexts (Motterlini 2007; Kanu and Leffler 2011; Parfenova et al. 2012b) and when induced at the right timing after stroke, vasodilation can be crucial in protecting the brain as it allows the supply of energy and nutrients which are in need in the ischemic tissue. The increase of regional blood flow due to cerebral vasodilation was previously found to be protective in stroke (Levi et al. 2012; Ginsberg 2016). The enhancement in the reperfusion of the lesioned tissue can be decisive for the final outcome after focal brain ischemia, and CO may be a good candidate therapeutic strategy acting at this level.

Leakage of the BBB was recently observed as early as 30 min after stroke (Shi et al. 2016). However, small and large molecules (≤150 kDa) only cross the BBB 3 h and 6 h after tMCAo. Moreover, permeability to very-large molecules (2000 kDa) was not detected until 24 h after injury (Shi et al. 2016). Furthermore, the infarct was found to expand from the striatum to the cerebral cortex within 6 h to 24 h, being the striatum the first brain region to suffer from BBB dysfunction (Shi et al. 2016). This body of evidence helps to understand the impact of CORM-A1 administration at 6 h after cerebral ischemia, and may explain the strong effects of CO in protecting the BBB. We might hypothesize that minimizing the BBB leakage to molecules with a size close to 2000 kDa is crucial for a better recovery, and CO may act on this mechanism. Further research is required to characterize whether CO can in fact prevent the stroke-induced increase in the permeability of the BBB to very-large molecules. The fact that CO fully protected the BBB, in contrast with the partial protection of the brain tissue under the conditions of this study, suggests that the vascular system and/or the glio-vascular unit is more responsive to CO treatment.

Functional preservation, rather than histological assessment, is presently considered the best approach to evaluate the outcome of the ischemic injury and the effect of putative protective strategies (Hattori et al. 2000). In this work, we evaluated the metabolic status of the cortical and striatal regions as an indirect measure of functionality after ischemic injury. At day 1 after MCAo we observed that CO administration promoted the maintenance of the metabolic load in the cerebral cortex, which is in accordance with the MRI results showing a reduced infarct area under the same conditions, although the results were not statistically significant in the latter experiments. At day 7 *post* injury, metabolic and histologic data were also in accordance since no protection was observed when the metabolite load was analyzed, and both cohorts presented a similar infarct volume. The time-dependence of the protective effect of CO is further reinforced by the evidence regarding the effect of CORM-A1 administration on the survival rate. A low rate of mortality was observed in mice subjected to MCAo while the CO-mediated responses were active. From day 3 onwards, when CORM-A1 was no longer administered, the difference between the two mice cohorts decreased, and at day 7 the survival rates were equal.

Quantification of the infarct volume using MRI suggests that the effects of CO are restricted to the cerebral cortex and limited to the early period after tMCAo. Region-specific effects of CO were previously described in the brain, either with low or high doses of CO (Nabeshima et al. 1991; Muraoka et al. 1998; Taskiran et al. 2007). However, the mechanisms underlying the differences in the regional responses to CO in the brain (*e*.*g*. cerebral cortex vs striatum) remain to be elucidated. The cortical lesion accounted for more than half of the total lesion under the conditions used in the present work. Therefore, the preferential effects in the cortical region are also reflected in the results obtained when the total lesion volume was analyzed. The fact that there is no significant lesion reduction when the BBB protection was so robust, points out for the need of further studies regarding the effect of CO on stroke.

In conclusion, the present work provides an integrated assessment of the time course of brain tissue and vasculature responses to CO in mice subjected to transient MCAo. Metabolic homeostasis was analyzed to characterize the effects of CO from the functional point of view. Damaged tissue responded to CO in a spatiotemporal dependent manner, which highly correlated with the effects in minimizing the loss in metabolic load. Moreover, the results suggest that a strong protection of the BBB and a slight metabolic preservation might contribute to effects of CO in re-establishing the brain and neuronal network and functional recovery after stroke. Causal effects of CO-mediated protection of the BBB on the metabolic stabilization needs to be address in the future. It is also important to determine whether CO prevents the infarct expansion and BBB extravasation from striatum to the cerebral cortex. With the first CO dose administered at 6h *post*-stroke, the present study shows that this strategy has a strong therapeutic potential for later treatments and ultimately may constitute an alternative for those patients that are not qualified for tPA administration. Longer CO treatments might overcome the loss of protection observed at 5 days after the last CO dose. In addition, CO administration may also be a good therapeutic option to minimize the risk of hemorrhagic transformation after tPA treatment, which is a major concern in clinical practice (Zhang et al. 2016).

## ACKNOWLEDGEMENTS

We thank Marius Widerøe for all the expertise in ^1^H-MRI data processing and analysis, as well as the fruitful discussions regarding ischemia. SRO was supported by a fellowship from Fundação para a Ciência e a Tecnologia (FCT) with reference SFRH/BD/51969/2012. This work was supported by FEDER (QREN) through Programa Mais Centro, under projects CENTRO-07-ST24-FEDER-002002, CENTRO-07-ST24-FEDER-002006 and CENTRO-07-ST24-FEDER-002008, through Programa Operacional Factores de Competitividade - COMPETE and national funds via FCT under projects Pest-C/SAU/LA0001/2013-2014, PTDC/SAU-NMC/120144/2010, PTDC/NEU-NMC/0198/2012 and FCT-ANR/NEU-NMC/0022/2012.

## SUPPLEMENTARY DATA

**Figure S1.**
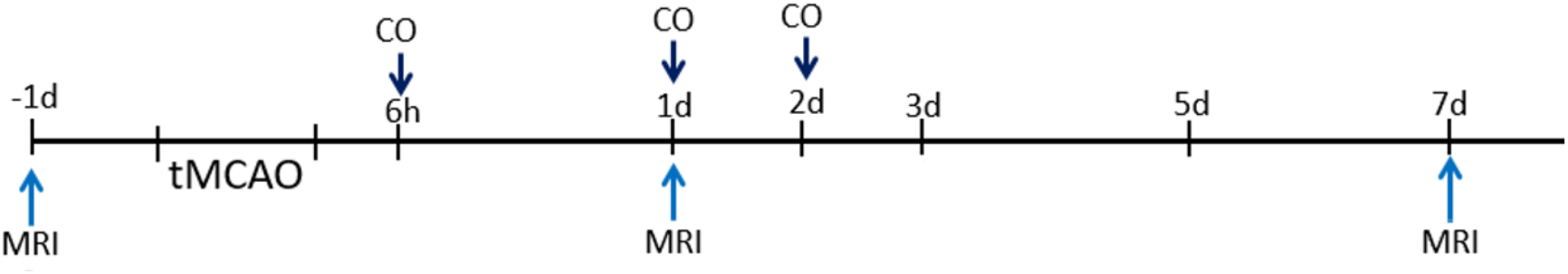
Schematic representation of the experimental design. Doses of CO (3 mg / kg) were given i.p. at 6 h, 1 day and 2 days after injury. Baseline MRI data was collected. MRI was performed at day 1 and day 7 *post*-tMCAo. Animals were euthanized at 7 days *post*-tMCAo. Brains were collected and analyzed, as described in the methods section.

**Figure S2.**
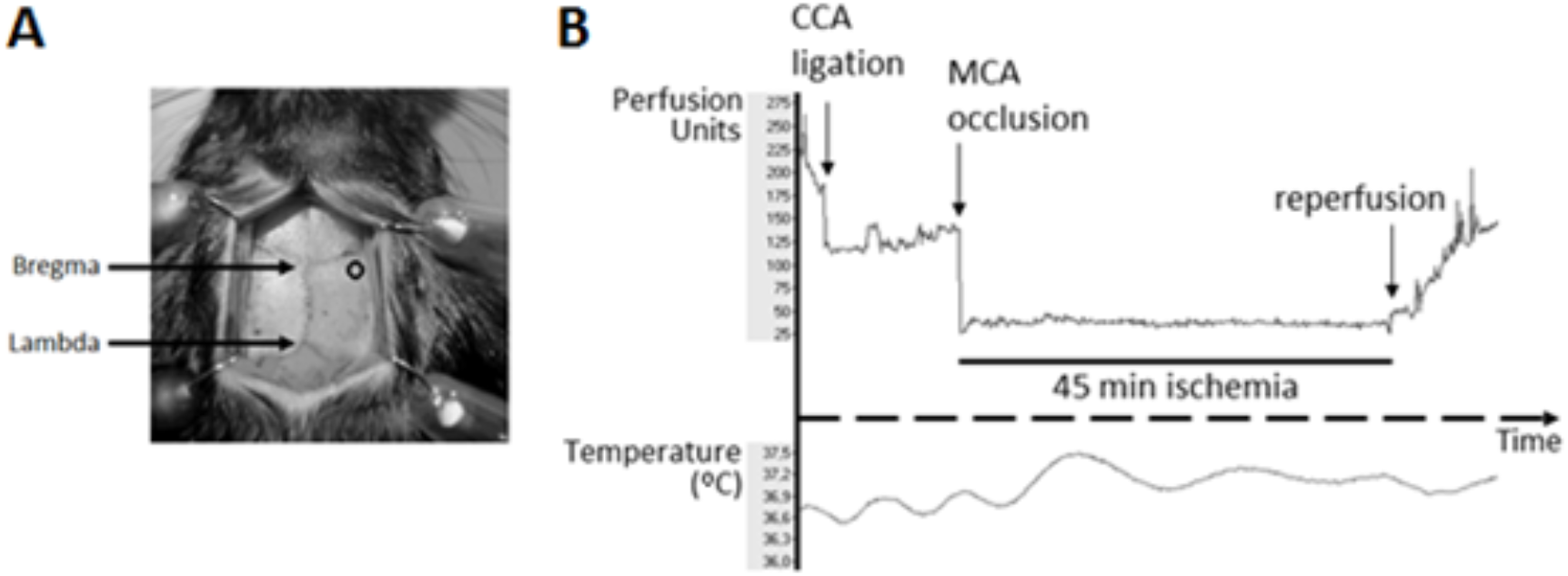
Laser Doppler microtip placement and representative profile of laser doppler flux and changes in body temperature in a tMCAo experiment. (A) Illustration of the location where the microtip was glued (open circle), approximately 2 mm posterior and 6 mm lateral to the bregma. (B) Laser-doppler flux profile measured over the right lateral parietal cortex during 45 min of middle cerebral artery occlusion (MCAo). Body temperature was measured with a rectal probe.

## REFERENCES

Almeida AS, Queiroga CSF, Sousa MFQ, et al (2012) Carbon monoxide modulates apoptosis by reinforcing oxidative metabolism in astrocytes: Role of Bcl-2. J Biol Chem 287:10761–10770. doi: 10.1074/jbc.M111.306738

Almeida AS, Soares NL, Sequeira CO, et al (2018) Improvement of neuronal differentiation by carbon monoxide: Role of pentose phosphate pathway. Redox Biol 17:338–347. doi: 10.1016/j.redox.2018.05.004

Almeida AS, Soares NL, Vieira M, et al (2016) Carbon Monoxide Releasing Molecule-A1 (CORM-A1) Improves Neurogenesis: Increase of Neuronal Differentiation Yield by Preventing Cell Death. PLoS One 11:e0154781. doi: 10.1371/journal.pone.0154781

Berridge K, Whishaw I (1992) Cortex, striatum and cerebellum: control of serial order in a grooming sequence. Exp Brain Res 90:275–290. doi: 10.1007/BF00227239

Berthet C, Lei H, Gruetter R, Hirt L (2011) Early Predictive Biomarkers for Lesion After Transient Cerebral Ischemia. Stroke 42:799–805. doi: 10.1161/STROKEAHA.110.603647

Bouët V, Freret T, Toutain J, et al (2007) Sensorimotor and cognitive deficits after transient middle cerebral artery occlusion in the mouse. Exp Neurol 203:555–567. doi: 10.1016/j.expneurol.2006.09.006

Chi P-L, Lin C-C, Chen Y-W, et al (2014a) CO Induces Nrf2-Dependent Heme Oxygenase-1 Transcription by Cooperating with Sp1 and c-Jun in Rat Brain Astrocytes. Mol Neurobiol. doi: 10.1007/s12035-014-8869-4

Chi P-L, Liu C-J, Lee I-T, et al (2014b) HO-1 Induction by CO-RM2 Attenuates TNF- α -Induced Cytosolic Phospholipase A 2 Expression via Inhibition of PKC α -Dependent NADPH Oxidase/ROS and NF-κ B. Mediators Inflamm 2014:1–18. doi: 10.1155/2014/279171

Choi YK, Maki T, Mandeville ET, et al (2016a) Dual effects of carbon monoxide on pericytes and neurogenesis in traumatic brain injury. Nat Med. doi: 10.1038/nm.4188

Choi YK, Park JH, Baek Y-Y, et al (2016b) Carbon monoxide stimulates astrocytic mitochondrial biogenesis via L-type Ca 2+ channel-mediated PGC-1α/ERRα activation. Biochem Biophys Res Commun 479:297–304. doi: 10.1016/j.bbrc.2016.09.063

Csongradi E (2012) Role of Carbon Monoxide in Kidney Function: Is a little Carbon Monoxide Good for the Kidney? Curr Pharm Biotechnol 13:819–826. doi: 10.2174/138920112800399284

Figueiredo-Pereira C, Dias-Pedroso D, Soares NL, Vieira HLA (2020) CO-mediated cytoprotection is dependent on cell metabolism modulation. Redox Biol. 32

Ginsberg MD (2016) Expanding the concept of neuroprotection for acute ischemic stroke: The pivotal roles of reperfusion and the collateral circulation. Prog Neurobiol. doi: 10.1016/j.pneurobio.2016.09.002

Harris JL, Choi I-Y, Brooks WM (2015) Probing astrocyte metabolism in vivo: proton magnetic resonance spectroscopy in the injured and aging brain. Front Aging Neurosci 7:. doi: 10.3389/fnagi.2015.00202

Hattori K, Lee H, Hurn PD, et al (2000) Cognitive Deficits After Focal Cerebral Ischemia in Mice Editorial Comment. Stroke 31:1939–1944. doi: 10.1161/01.STR.31.8.1939

Hoffmann CJ, Harms U, Rex A, et al (2015) Vascular Signal Transducer and Activator of Transcription-3 Promotes Angiogenesis and Neuroplasticity Long-Term After Stroke. Circulation 131:1772–1782. doi: 10.1161/CIRCULATIONAHA.114.013003

Jiang Q, Ewing JR, Ding GL, et al (2005) Quantitative evaluation of BBB permeability after embolic stroke in rat using MRI. J Cereb Blood Flow Metab 25:583–592. doi: 10.1038/sj.jcbfm.9600053

Kalueff A V, Stewart AM, Song C, et al (2015) Neurobiology of rodent self-grooming and its value for translational neuroscience. Nat Rev Neurosci 17:45–59. doi: 10.1038/nrn.2015.8

Kanu A, Leffler CW (2011) Arachidonic acid- and prostaglandin E2-induced cerebral vasodilation is mediated by carbon monoxide, independent of reactive oxygen species in piglets. Am J Physiol Heart Circ Physiol 301:H2482–2487. doi: 10.1152/ajpheart.00628.2011

Kassner A, Merali Z (2015) Assessment of Blood–Brain Barrier Disruption in Stroke. Stroke 46:3310–3315. doi: 10.1161/STROKEAHA.115.008861

Kim KM, Pae HO, Zheng M, et al (2007) Carbon monoxide induces heme oxygenase-1 via activation of protein kinase R-like endoplasmic reticulum kinase and inhibits endothelial cell apoptosis triggered by endoplasmic reticulum stress. Circ Res 101:919–927. doi: 10.1161/CIRCRESAHA.107.154781

Klaus JA, Kibler KK, Abuchowski A, Koehler RC (2010) Early Treatment of Transient Focal Cerebral Ischemia with Bovine PEGylated Carboxy Hemoglobin Transfusion. Artif Cells, Blood Substitutes, Biotechnol 38:223–229. doi: 10.3109/10731199.2010.488635

Knecht KR, Milam S, Wilkinson D a, et al (2010) Time-dependent action of carbon monoxide on the newborn cerebrovascular circulation. Am J Physiol Heart Circ Physiol 299:H70–H75. doi: 10.1152/ajpheart.00258.2010

Krueger M, Bechmann I, Immig K, et al (2015) Blood-brain barrier breakdown involves four distinct stages of vascular damage in various models of experimental focal cerebral ischemia. J Cereb Blood Flow Metab 35:292–303. doi: 10.1038/jcbfm.2014.199

Leffler CW, Parfenova H, Jaggar JH (2011) Carbon monoxide as an endogenous vascular modulator. Am J Physiol - Hear Circ Physiol 301:H1–H11. doi: 10.1152/ajpheart.00230.2011

Lei H, Berthet C, Hirt L, Gruetter R (2009) Evolution of the neurochemical profile after transient focal cerebral ischemia in the mouse brain. J Cereb Blood Flow Metab 29:811–819. doi: 10.1038/jcbfm.2009.8

Levi H, Schoknecht K, Prager O, et al (2012) Stimulation of the Sphenopalatine Ganglion Induces Reperfusion and Blood-Brain Barrier Protection in the Photothrombotic Stroke Model. PLoS One 7:e39636. doi: 10.1371/journal.pone.0039636

Mahan VL (2012) Neuroprotective, neurotherapeutic, and neurometabolic effects of carbon monoxide. Med Gas Res 2:32. doi: 10.1186/2045-9912-2-32

Maines MD (2000) The heme oxygenase system and its functions in the brain. Cell Mol Biol (Noisy-le-grand) 46:573–85

McColl BW, Carswell H V., McCulloch J, Horsburgh K (2004) Extension of cerebral hypoperfusion and ischaemic pathology beyond MCA territory after intraluminal filament occlusion in C57Bl/6J mice. Brain Res 997:15–23. doi: 10.1016/j.brainres.2003.10.028

Meurer WJ, Barth BE, Gaddis G, et al (2016) Rapid Systematic Review: Intra-Arterial Thrombectomy (“Clot Retrieval”) for Selected Patients with Acute Ischemic Stroke. J Emerg Med. doi: 10.1016/j.jemermed.2016.10.004

Motterlini R (2007) Carbon monoxide-releasing molecules (CO-RMs): vasodilatory, anti-ischaemic and anti-inflammatory activities. Biochem Soc Trans 35:1142–6. doi: 10.1042/BST0351142

Motterlini R, Sawle P, Hammad J, et al (2005) CORM-A1: a new pharmacologically active carbon monoxide-releasing molecule. Faseb J 19:284–286. doi: 10.1096/fj.04-2169fje

Muraoka M, Hayakawa H, Kagaya A, et al (1998) Effects of carbon monoxide exposure on serotonergic neuronal systems in rat brain. Life Sci 62:2101–8

Nabeshima T, Katoh A, Ishimaru H, et al (1991) Carbon monoxide-induced delayed amnesia, delayed neuronal death and change in acetylcholine concentration in mice. J Pharmacol Exp Ther 256:378–84

Obeso JA, Lanciego JL (2011) Past, Present, and Future of the Pathophysiological Model of the Basal Ganglia. Front Neuroanat 5:. doi: 10.3389/fnana.2011.00039

Oliveira SR, Figueiredo-Pereira C, Duarte CB, Vieira HLA (2019) P2×7 Receptors Mediate CO-Induced Alterations in Gene Expression in Cultured Cortical Astrocytes—Transcriptomic Study. Mol Neurobiol 56:3159–3174. doi: 10.1007/s12035-018-1302-7

Parfenova H, Leffler CW, Basuroy S, et al (2012a) Antioxidant roles of heme oxygenase, carbon monoxide, and bilirubin in cerebral circulation during seizures. J Cereb Blood Flow Metab 32:1024–34. doi: 10.1038/jcbfm.2012.13

Parfenova H, Tcheranova D, Basuroy S, et al (2012b) Functional role of astrocyte glutamate receptors and carbon monoxide in cerebral vasodilation response to glutamate. AJP Hear Circ Physiol 302:H2257–H2266. doi: 10.1152/ajpheart.01011.2011

Park S-Y, Marasini S, Kim G-H, et al (2014) A Method for Generate a Mouse Model of Stroke: Evaluation of Parameters for Blood Flow, Behavior, and Survival. Exp Neurobiol 23:104. doi: 10.5607/en.2014.23.1.104

Prinz V, Köning J, Ji S, et al (2010) Standard operating procedures (SOP) in experimental stroke research: SOP for middle cerebral artery occlusion in the mouse. Nat Preced 202213:1–5. doi: 10.1038/npre.2010.3492.2

Queiroga CSF, Alves RMA, Conde S V., et al (2016) Paracrine effect of carbon monoxide –astrocytes promote neuroprotection through purinergic signaling in mice. J Cell Sci 129:3178–3188. doi: 10.1242/jcs.187260

Queiroga CSF, Tomasi S, Widerøe M, et al (2012) Preconditioning Triggered by Carbon Monoxide (CO) Provides Neuronal Protection Following Perinatal Hypoxia-Ischemia. PLoS One 7:. doi: 10.1371/journal.pone.0042632

Queiroga CSF, Vercelli A, Vieira HLA (2015) Carbon monoxide and the CNS: challenges and achievements. Br J Pharmacol 172:1533–1545. doi: 10.1111/bph.12729

Ryter SW, Choi AMK (2009) Heme oxygenase-1/carbon monoxide: From metabolism to molecular therapy. Am J Respir Cell Mol Biol 41:251–260. doi: 10.1165/rcmb.2009-0170TR

Saransaari P, Oja SS (2010) Modulation of taurine release in ischemia by glutamate receptors in mouse brain stem slices. Amino Acids 38:739–746. doi: 10.1007/s00726-009-0278-z

Schwamm LH, Ali SF, Reeves MJ, et al (2013) Temporal Trends in Patient Characteristics and Treatment With Intravenous Thrombolysis Among Acute Ischemic Stroke Patients at Get With the Guidelines-Stroke Hospitals. Circ Cardiovasc Qual Outcomes 6:543–549. doi: 10.1161/CIRCOUTCOMES.111.000303

Shi Y, Zhang L, Pu H, et al (2016) Rapid endothelial cytoskeletal reorganization enables early blood–brain barrier disruption and long-term ischaemic reperfusion brain injury. Nat Commun 7:10523. doi: 10.1038/ncomms10523

Taskiran D, Nesil T, Alkan K (2007) Mitochondrial oxidative stress in female and male rat brain after ex vivo carbon monoxide treatment. Hum & Exp Toxicol 26:645–651. doi: 10.1177/0960327107076882

Wang B, Cao W, Biswal S, Doré S (2011) Carbon monoxide-activated Nrf2 pathway leads to protection against permanent focal cerebral ischemia. Stroke 42:2605–2610. doi: 10.1161/STROKEAHA.110.607101

Wardlaw JM, Murray V, Berge E, del Zoppo GJ (2014) Thrombolysis for acute ischaemic stroke. In: Wardlaw JM (ed) Cochrane Database of Systematic Reviews. John Wiley & Sons, Ltd, Chichester, UK, pp 1581–1588

Winter B, Juckel G, Viktorov I, et al (2005) Anxious and Hyperactive Phenotype Following Brief Ischemic Episodes in Mice. Biol Psychiatry 57:1166–1175. doi: 10.1016/j.biopsych.2005.02.010

Wunder A, Schoknecht K, Stanimirovic DB, et al (2012) Imaging blood-brain barrier dysfunction in animal disease models. Epilepsia 53 Suppl 6:14–21. doi: 10.1111/j.1528-1167.2012.03698.x

Yabluchanskiy A, Sawle P, Homer-Vanniasinkam S, et al (2012) CORM-3, a carbon monoxide-releasing molecule, alters the inflammatory response and reduces brain damage in a rat model of hemorrhagic stroke. Crit Care Med 40:544–552. doi: 10.1097/CCM.0b013e31822f0d64

Zeynalov E, Dore S (2009) Low doses of carbon monoxide protect against experimental focal brain ischemia. Neurotox Res 15:133–137. doi: 10.1007/s12640-009-9014-4

Zhang J, Ho W, Reis C, et al (2016) Pharmacological Management Options to Prevent and Reduce Ischemic Hemorrhagic Transformation. Curr Drug Targets 17:1–1. doi: 10.2174/1389450117666160818115850

